# Systematic characterization of indel variants using a yeast-based protein folding sensor

**DOI:** 10.1101/2024.07.11.603017

**Authors:** Sven Larsen-Ledet, Søren Lindemose, Aleksandra Panfilova, Sarah Gersing, Caroline H. Suhr, Aitana Victoria Genzor, Heleen Lanters, Sofie V. Nielsen, Kresten Lindorff-Larsen, Jakob R. Winther, Amelie Stein, Rasmus Hartmann-Petersen

## Abstract

Gene variants resulting in insertions or deletions of amino acid residues (indels) have important consequences for evolution and are often linked to disease, yet compared to missense variants the effects of indels are poorly understood and predicted. To approach this issue, we developed a sensitive protein folding sensor based on complementation of uracil auxotrophy in yeast by circular permutated orotate phosphoribosyltransferase (CPOP). The sensor accurately reports on the folding of disease-linked missense variants and *de novo* designed proteins. Applying the folding sensor to a saturated library of single amino acid indel variants in human DHFR revealed that most regions which tolerate indels are confined to internal loops and the N- and C-termini. Surprisingly, indels are also allowed at a central α-helix. Several indels are temperature-sensitive and the folding of most of these indels is rescued upon binding to the competitive DHFR inhibitor methotrexate. Rosetta and AlphaFold2 predictions correlate with the observed effects, suggesting that most indels operate by destabilizing the native fold and that these computational tools may be useful for classification of indels observed in population sequencing.

## Introduction

The folding, stability and function of a protein are determined by its amino acid sequence [1]. Accordingly, gene variants that affect the amino acid sequence can result in proteins with changed structural, biophysical and functional properties [2]. In recent years, tremendous progress has been made in our understanding and ability to predict the consequences of single amino acid substitutions (missense variants) on the structural stability and function of proteins [2–7]. In comparison, the consequences of amino acid insertions and deletions (indels) are not well understood and poorly predicted [8, 9]. This is despite indels being abundant in human genomes [10–12], often being connected with inherited traits and genetic disorders [10, 11, 13], and likely to be valuable in protein engineering [14–16].

Indels are fundamentally distinct from missense variants in that they not only alter the amino acid sidechains, but also affect the backbone length. Accordingly, the consequences of indel variants are often more severe. Thus, indels are subject to stronger purifying selection than missense variants during evolution [17] and are typically enriched in intrinsically disordered regions (IDRs) [12]. Within structured regions, a single amino acid insertion or deletion will cause a 100° rotation and 1.5 Å translation in an α-helix, and a 180° rotation and 3.8 Å translation in a β-strand [14]. Nonetheless, many proteins can still accommodate indels even within structured regions by local adaptation of the fold [14–16]. Albeit indels often result in loss-of-function (LOF) phenotypes, as exemplified by the CFTR F508Δ variant linked to cystic fibrosis [18], some indels may be beneficial, as for instance the EGFP G4Δ variant that displays enhanced folding and fluorescence properties [19].

Recently, a few reports have systematically analyzed the effects of indels in selected proteins by so-called deep mutational scanning (DMS) or multiplexed assays of variant effects (MAVEs) technologies [20–27]. The results of these efforts seem to converge on the following rules: Typically, indels appear to be tolerated best in IDRs, flexible loops as well as the N- and C-terminal regions. Within structured regions, indels are better tolerated in α-helices and at the ends of β-strands, and least tolerated within the central parts of a β-strand (reviewed in [16]). In addition, it seems that sites that tolerate insertions tend to overlap with sites that tolerate deletions [23, 24, 26]. It further appears that the site of insertion is more important than the number (1-3 residues) [24] or nature of amino acids inserted [26, 27]. However, further research on a more diverse set of proteins is needed to confirm this.

Unlike missense variants that tend to be allowed at surface-exposed positions [28], this is not always the case for indels [26, 27], and indel tolerance appears to vary widely both between proteins and within different regions of a protein [27]. Accordingly, current computational variant effect predictors are either not very accurate or do not provide scores for indels [29]. Here, we systematically characterize single amino acid indels in the human dihydrofolate reductase (DHFR) using a protein folding sensor constructed on a circular permutated variant of orotate phosphoribosyltransferase (CPOP) [30]. In line with previous studies, we find that most regions that tolerate indels are confined to internal loops and the N- and C-termini. Overall, deletions appear to be more detrimental than insertions. Surprisingly, we also find loops that do not tolerate indels as well as a central α-helix where indels are tolerated. Furthermore, compromised folding of most indel variants is rescued at low temperatures or in the presence of the competitive DHFR inhibitor methotrexate (MTX). We find that Rosetta and AlphaFold2 (AF2) predictions correlate with the observed effects, suggesting these tools may hold promise for providing a mechanistic interpretation and making genotype-phenotype predictions for indels in structured proteins.

## Results

### A yeast-based protein folding sensor

To assess protein folding and stability in yeast cells, we and others have previously directly fused the protein of interest (POI) to GFP and/or Ura3, thus monitoring the folding and abundance of the POI by GFP fluorescence or complementation of uracil auxotrophy, respectively [31–33]. However, fusing a POI to a different folded protein such as GFP and Ura3 may modulate its stability [34] and change the dynamic range of a folding sensor.

As an alternative, we used a different approach based on a circular permutated orotate phosphoribosyltransferase (CPOP) which has previously proven useful as a protein folding sensor in *E. coli* [30]. In this system, *E. coli* orotate phosphoribosyltransferase (OPRTase) is circular permutated, so the N-terminal region encompassing residues 1-92 is moved to the C-terminus at position 213 (**Fig. 1A**). The resulting circular permutated OPRTase (CPOP) is active as an enzyme [30], but as typical for circular permutated proteins [35], displays reduced structural stability [30], and is therefore thought not to perturb substantially the stability of a POI inserted into a linker between the original C- and N-termini (**Fig. 1A**). However, due to its positioning between the C- and N-terminal halves, even slight changes in the structure of the POI should compromise CPOP folding and result in insufficient OPRTase activity to completement the deletion of the yeast OPRTase genes *URA5* and *URA10* (**Fig. 1B**). OPRTase functions as a homodimer with two active sites situated at the interface of the monomers (**Fig. 1C**). The fact that OPRTase is a dimer likely increases the sensitivity of the CPOP folding sensor and makes the folding/unfolding of the fusion protein more cooperative. To adapt the CPOP system to yeast, we first compared the growth of wild-type *S. cerevisiae* with strains deleted for the OPRTase orthologues *URA5* and *URA10*. In agreement with previous reports [36, 37], the strains appeared wild type in presence of uracil. While there was no obvious uracil-dependent growth defect of the *ura10*Δ strain [37], the *ura5*Δ strain displayed a strong growth defect in the absence of uracil [36], and a *ura5*Δ*ura10*Δ double mutant appeared completely auxotroph for uracil (**Fig. 2A**) [37]. In addition, we noted that none of the strains were temperature-sensitive (**Fig. 2A**).

**Fig. 1.**
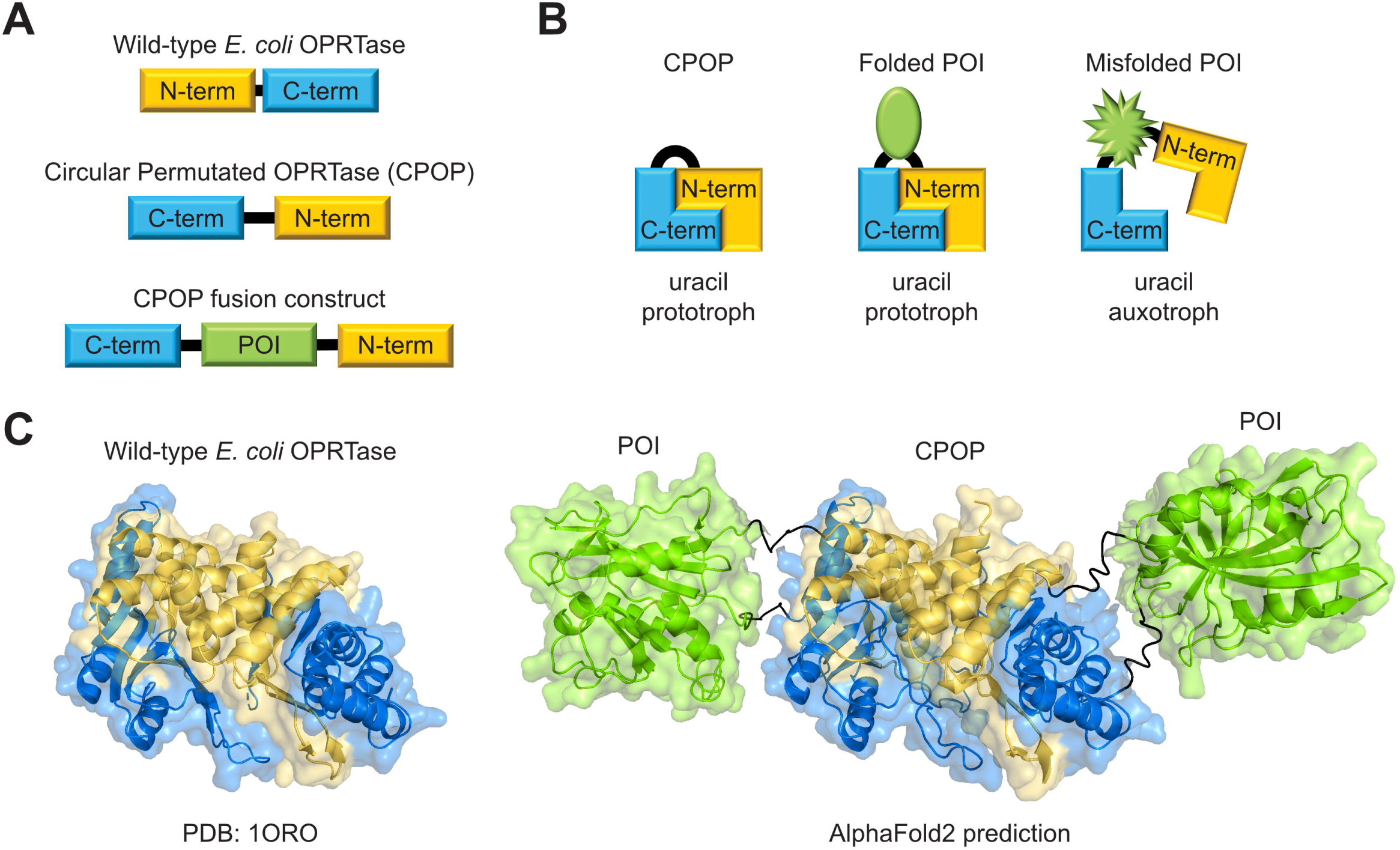
Construction of circular permutated orotate phosphoribosyltransferase (CPOP). (A) Wild-type *E. coli* OPRTase was circularly permutated by inserting a start codon before residue 92 and a stop codon after residue 91. This resulted in the rearrangement of the N- and C-termini, which were separated by a short linker, leading to the construction of CPOP. A CPOP fusion construct was created by introducing a protein of interest (POI) in the short linker between the original C- and N-termini. (B) CPOP is an active, but structurally compromised enzyme and its folding depends on the stability of the POI. Misfolding of the POI results in decreased CPOP activity and renders yeast cells uracil auxotrophic when lacking endogenous OPRTase activity. (C) The crystal structure of wild-type *E. coli* OPRTase (PDB: 1ORO) shown as the homodimer with the N-terminal region shown in yellow and C-terminal region shown in blue (left). A POI (here human DHFR) is inserted into the linker of CPOP to construct CPOP-DHFR (right). AF2 was used to predict the structure of the monomeric CPOP-DHFR fusion construct, which was duplicated and aligned to each of the monomers of 1ORO. The N- and C-termini are colored as in 1ORO, DHFR is shown in green, and the linkers are shown in black.

**Fig. 2.**
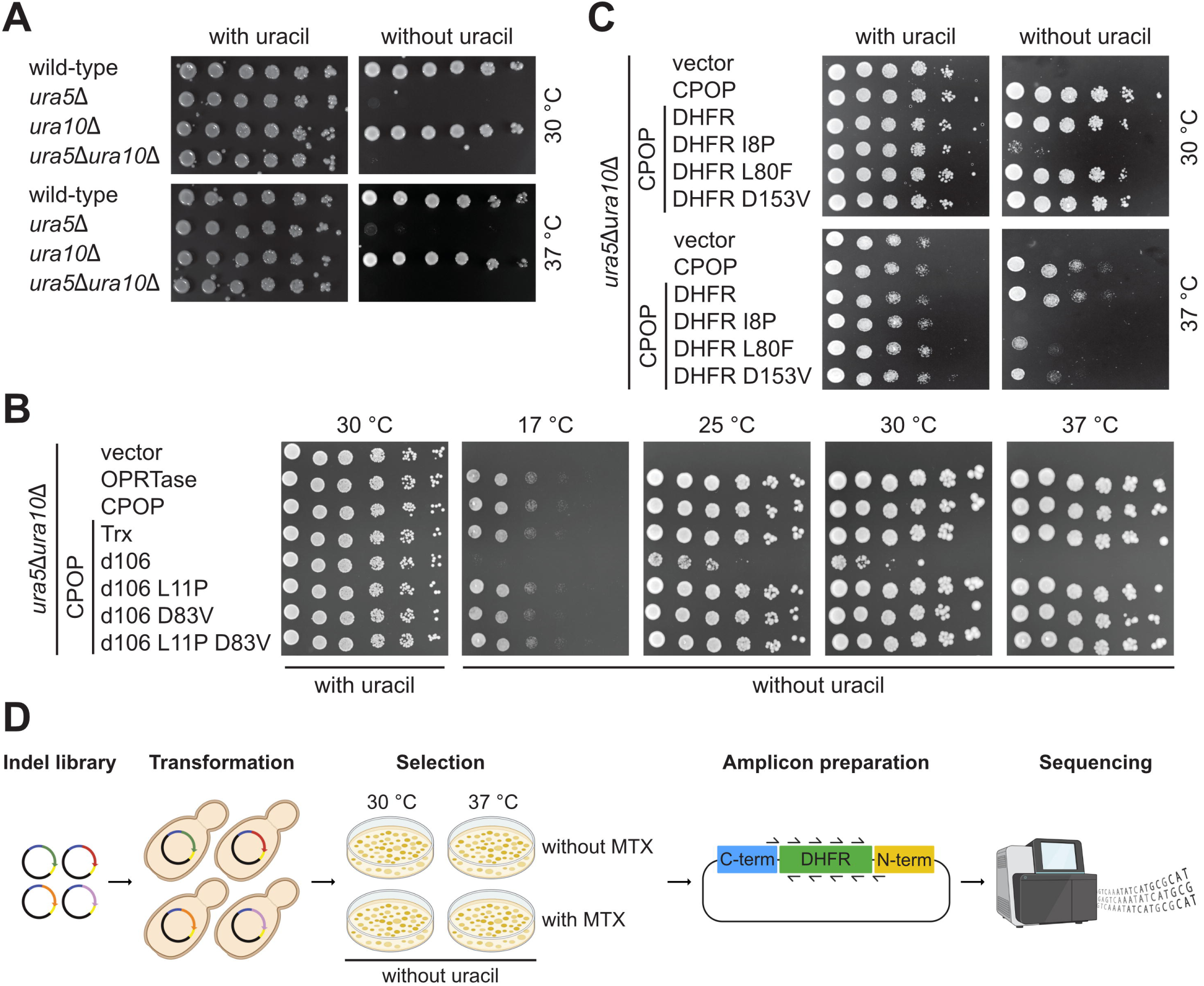
CPOP functions as a folding sensor in the ura5Δura10Δ strain. (A) Yeast growth assays of wild-type, *ura5*Δ, *ura10*Δ and *ura5*Δ*ura10*Δ yeast strains on medium with and without uracil at 30 °C and 37 °C. (B) Comparison of growth of the *ura5*Δ*ura10*Δ strain transformed with either an empty vector, wild-type *E. coli* OPRTase, CPOP or CPOP fusion constructs with wild-type *E. coli* Trx, the *de novo* Trx design dF106 or stabilized dF106 variants. Cells were grown on medium with and without uracil at 17 °C, 25 °C, 30 °C and 37 °C. (C) Comparison of growth of the *ura5*Δ*ura10*Δ strain transformed with either an empty vector, CPOP or CPOP fusion constructs with the indicated DHFR variants. Cells were grown on medium with and without uracil at 30 °C and 37 °C. (D) Schematic overview of the experimental design to screen DHFR indel libraries in the CPOP folding sensor. The libraries comprised the wild-type sequence, a synonymous wild-type variant at each position, a stop codon (nonsense mutation) at each position, all single amino acid deletions and a single glycine insertion after each residue. Panel (D) was created with BioRender.com.

Next, we continued to test if *E. coli* OPRTase and CPOP were able to complement the uracil auxotrophy of the *ura5*Δ*ura10*Δ strain. The genes encoding OPRTase and CPOP were expressed from the constitutive *GAP* promoter in a centromere-based plasmid (**Supplementary Fig. 1**). Indeed, both OPRTase and CPOP complemented the uracil auxotrophy over a range of temperatures (**Fig. 2B**), thus allowing us to monitor folding of a POI inserted into the linker in CPOP under these conditions.

Previously, CPOP has been used in *E. coli* to assess the folding and stability of *E. coli* thioredoxin (Trx) and various artificial thioredoxins generated by *de novo* protein design, including the dF106 protein and various dF106 point mutants with increased stability [30]. To further test the sensitivity of the yeast-based CPOP system, we compared the growth of cells expressing CPOP carrying Trx, dF106 and three dF106 point mutants with increased stability (L11P, D83V, and L11P D83V double mutant). In line with the results from *E. coli* [30], cells with the artificially designed dF106 protein in the CPOP sensor displayed poor growth, while the stabilized dF106 point mutants appeared similar to wild-type Trx (**Fig. 2B**).

In conclusion, this suggests that the yeast-based CPOP system reflects the previous results in *E. coli* and thus allows coupling of protein folding *in vivo* to cell growth.

### CPOP-mediated detection of disease-linked missense variants with protein folding defects

Reduced structural stability of the encoded protein is a well-characterized mechanism by which missense mutations can result in loss-of-function phenotypes [2]. We have previously shown this to be the case for missense variants in the human DHFR enzyme linked to recessive megaloblastic anemia (OMIM: 613839) [31]. Specifically, the pathogenic DHFR variants L80F and D153V are both slightly structurally destabilized [31]. To test if the CPOP sensor was sensitive enough to allow detection of the disease-linked DHFR variants, we inserted wild-type human DHFR and the L80F and D153V variants into the CPOP folding sensor. In addition, we included the I8P variant, which has not been observed in patients, but which we have predicted [38] would lead to a more dramatic loss of protein stability.

Indeed, the I8P variant led to strongly reduced growth in the absence of uracil at 30 °C and complete lack of growth at 37 °C (**Fig. 2C**). In comparison, the disease-linked L80F and D153V variants appeared stable at 30 °C, but displayed slow growth at 37 °C, indicating that these protein variants are indeed poorly folded, albeit not to the same extent as I8P.

Based on these results, we conclude that the CPOP folding sensor is sensitive to even minor changes in protein folding and stability.

### Systematic analyses of DHFR indel variants using the CPOP folding sensor

Next, we decided to apply the CPOP folding sensor in a screen aimed at characterizing the effects of indel variants in human DHFR. We selected DHFR, as this is a short (187 residues) and well-characterized enzyme, which is structurally stabilized upon binding to its inhibitor methotrexate (MTX) [31, 39].

We divided the DHFR sequence into five tiles overlapping by three amino acid residues (9 bp) and designed, screened, and sequenced libraries separately in each tile. The DHFR libraries comprised all possible single amino acid deletions as well as a single glycine insertion after each residue. In addition, we included the wild-type sequence as well as a synonymous wild-type (silent) variant and a stop codon (nonsense mutation) at each position. The DHFR indel libraries were inserted into the CPOP linker and transformed into the *ura5*Δ*ura10*Δ yeast strain. From a dense transformation culture, some cells were harvested to serve as the control condition prior to selection. For selection of protein folding, cells were plated on medium without uracil in triplicates. To identify indels with temperature-sensitive folding defects the plated cells were incubated at both 30 °C and 37 °C. In addition, we also included plates containing MTX to determine if any of the DHFR indels were stabilized upon binding to MTX. After two days incubation, the cells were washed off the plates and used for plasmid purification, amplicon preparation and paired-end Illumina sequencing (**Fig. 2D**). We processed the sequencing reads using an in-house pipeline (see Materials and Methods) to call and quantify single amino acid indels, synonymous mutations, and nonsense mutations. The frequency of each variant relative to the synonymous variants in the control condition and on the selection plates allowed us to determine a CPOP folding score per variant at four different conditions (30 °C, 30 °C + MTX, 37 °C and 37 °C + MTX). The CPOP scores were scaled to range from 0 (folding of nonsense variants) to 1 (folding of wild-type and synonymous variants) independently at each experimental condition.

The obtained dataset contains CPOP scores for at least one condition for 186 out of 187 (99%) possible insertions, 186 out of 187 (99%) possible deletions, 176 out of 177 (99%) possible synonymous wild-type variants and 185 out of 186 (99%) possible nonsense variants. In all conditions, the read counts showed high correlation between the three replicates (**Supplementary Fig. 2**) and the CPOP scores for DHFR indel variants displayed low standard deviations (**Supplementary Fig. 3**). In all conditions, the nonsense variants displayed folding scores around 0 and the synonymous variants displayed scores around 1 (**Fig. 3A, Supplementary Table 1**). The distribution of the indel variants largely appeared bimodal, with the majority of the variants clustering around either 0 or 1, while the remainder displayed intermediate scores (**Fig. 3A**). For validation, we analyzed the growth of wild-type DHFR along with five different insertion and five different deletion variants in low throughput. The results from these experiments matched those obtained by the screen (**Supplementary Fig. 4**).

**Fig. 3.**
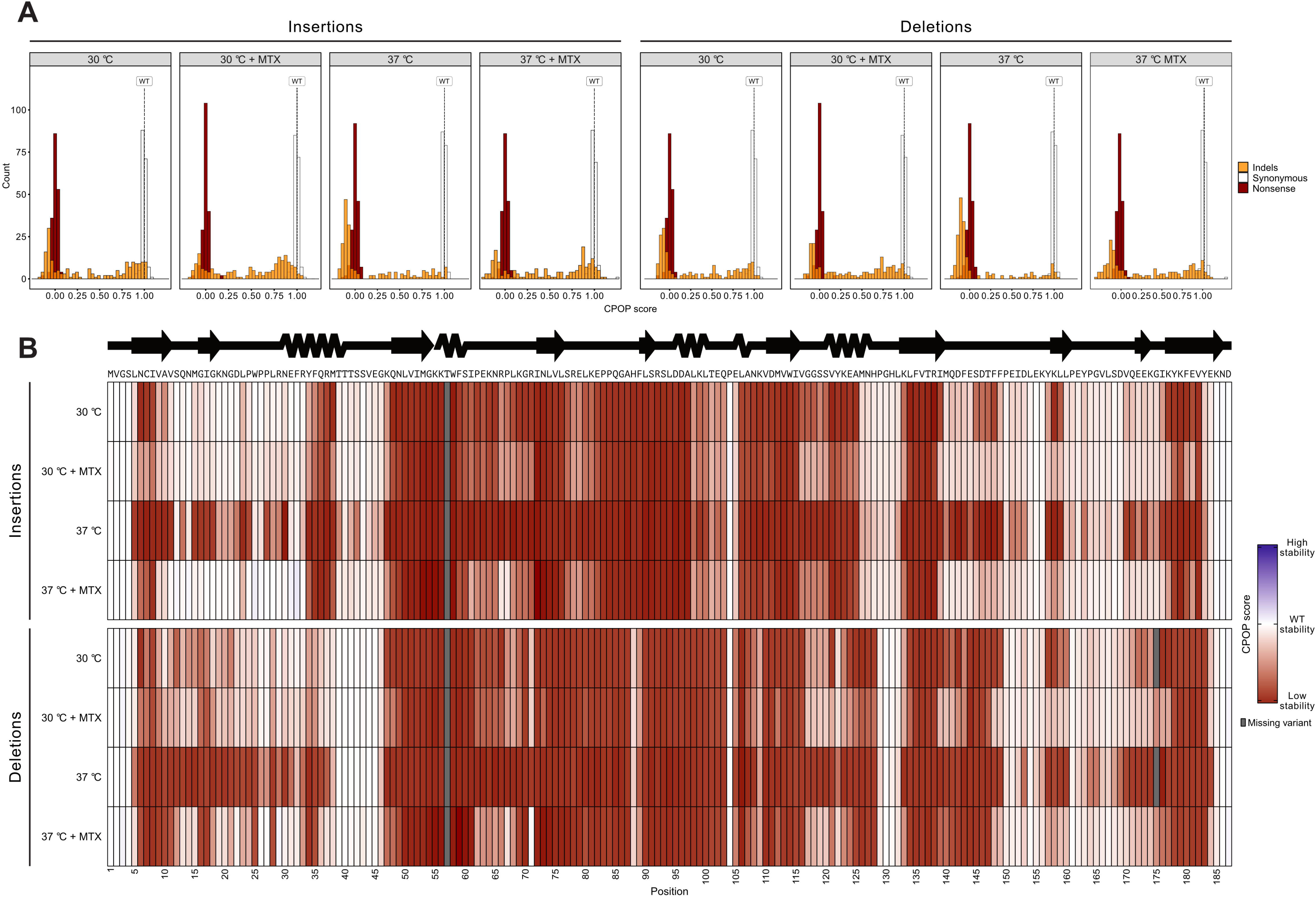
Results of the DHFR indel screen using the CPOP folding sensor. (A) CPOP score distributions for insertions and deletions (orange), synonymous variants (white) and nonsense variants (red) at the four different conditions (30 °C, 30 °C + MTX, 37 °C and 37 °C + MTX). The CPOP scores were rescaled, so a score of 0 corresponds to complete loss of stability (nonsense variants) and a score of 1 corresponds to wild-type-like stability (synonymous variants). A score above 1 corresponds to increased stability. (B) Heatmaps of CPOP scores for insertions and deletions at the four different conditions for each position in DHFR. The CPOP scores range from low stability (red) over wild-type-like stability (white) to high stability (blue). Missing variants are colored grey. The amino acid sequence and a linear representation of the secondary structure of human DHFR are shown above the heatmap.

As expected, many indel variants were detrimental to DHFR folding, and at most positions the deletions appeared more detrimental than insertions of glycine (**Fig. 3B, Supplementary Fig. 5**). However, in general the positions that were sensitive to insertions were also sensitive to deletions. For the α-helix spanning residues 30-40, insertions were not tolerated, while deletions were (**Fig. 3B**). For both insertions and deletions, the detrimental effects were exacerbated at 37 °C and the otherwise tolerant region spanning residues 10-30 became highly sensitive (**Fig. 3B**). In the presence of MTX, the sensitive positions towards the N- and C-termini were strongly stabilized (**Fig. 3B**). These effects were also evident when we mapped the indel scores onto the DHFR structure, and moreover revealed that the ends of the β-strands appeared to tolerate indels better than the central parts (**Fig. 4A**). In addition, deletions were more detrimental in β-strands than in α-helices, which was not the case for insertions (**Fig. 4B**). Not surprisingly, α-helices and β-strands were more sensitive to indels than loops at most conditions (**Fig. 4B**). However, we noticed that the central loops spanning residues 60-70 and 75-88 were highly sensitive to indels, despite being long and exposed (**Fig. 3B**, **Fig. 4A**). This observation exemplifies that backbone perturbations by indels can have surprising effects in regions where missense mutations are expected to be tolerated. Consistent with this, indel tolerance did not correlate with relative solvent accessible surface area (rSASA) (**Supplementary Fig. 6**).

**Fig. 4.**
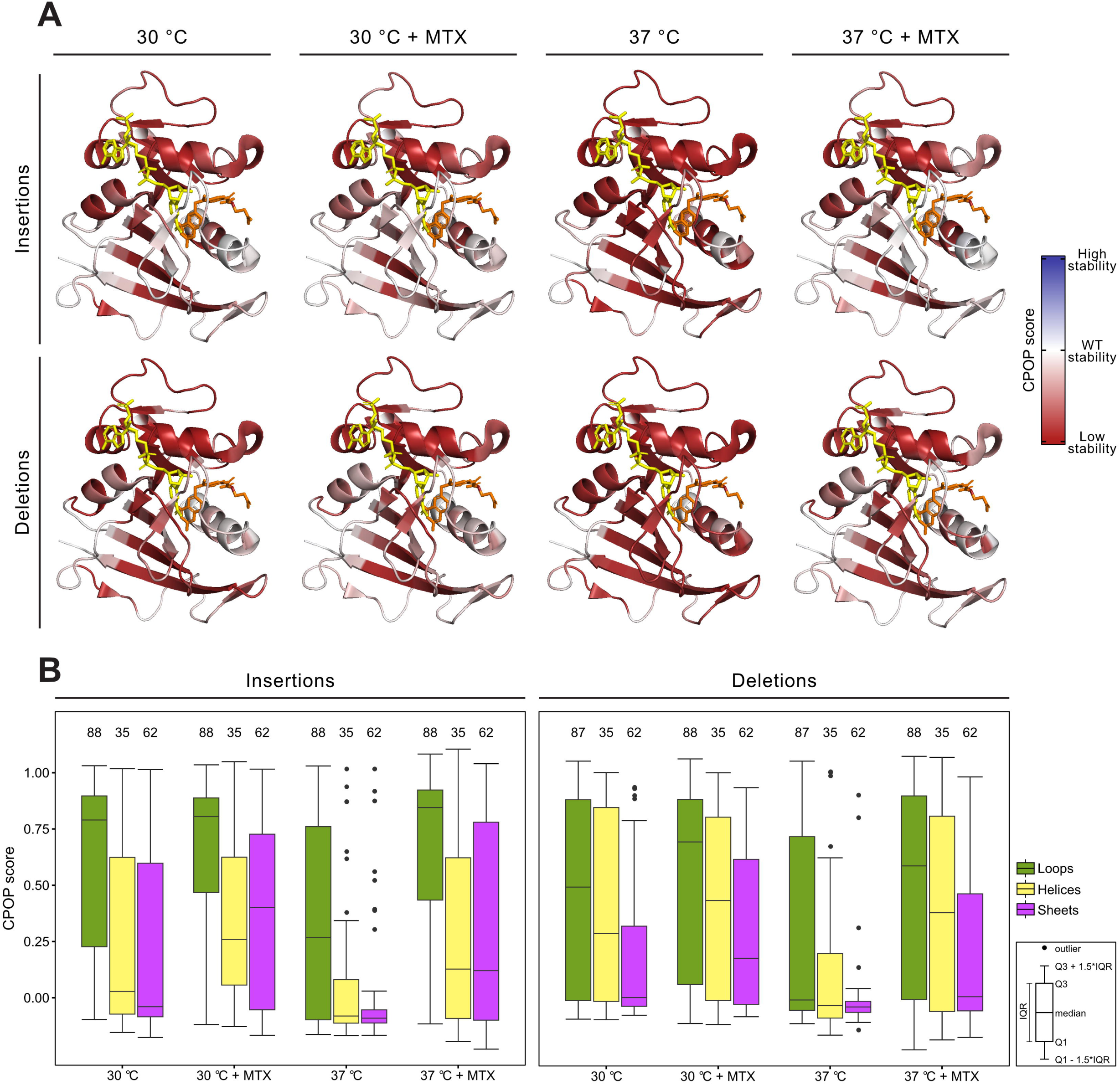
Structural effects of DHFR insertions and deletions. (A) The CPOP scores for insertions and deletions at the four different conditions are mapped to the DHFR structure (PDB: 1U72). The color scheme is the same as in Fig. 3B. NADPH is shown in yellow and MTX is shown in orange. (B) Box plots illustrating the CPOP scores for insertions and deletions at the four different conditions for positions in loops, helices, and strands. The three secondary structure elements were assigned for DHFR (PDB: 1U72) using the DSSP algorithm. The number of positions included is indicated above each box plot. The black dots indicate outliers. The box plots are defined as indicated in the legend.

To better visualize positions that were critical for temperature sensitivity or rescue by MTX, we determined the differences in the scores, generating ΔTemperature and ΔMTX scores (**Fig. 5A**) and mapped these onto the DHFR structure (**Fig. 5B**). This confirmed the trends described above and revealed that many of the temperature-sensitive and MTX-dependent positions overlapped (**Fig. 5B**). While some of the MTX-dependent positions were situated near the active site (where MTX binds), many were located surprisingly far away (**Fig. 5B**). Thus, the ΔMTX scores did not correlate with distance to the active site (**Supplementary Fig. 7**). This agrees with *in vitro* and *in vivo* ligand-binding studies on *E. coli* DHFR [39–42] and suggests that temperature and MTX-binding globally affect the protein conformation.

**Fig. 5.**
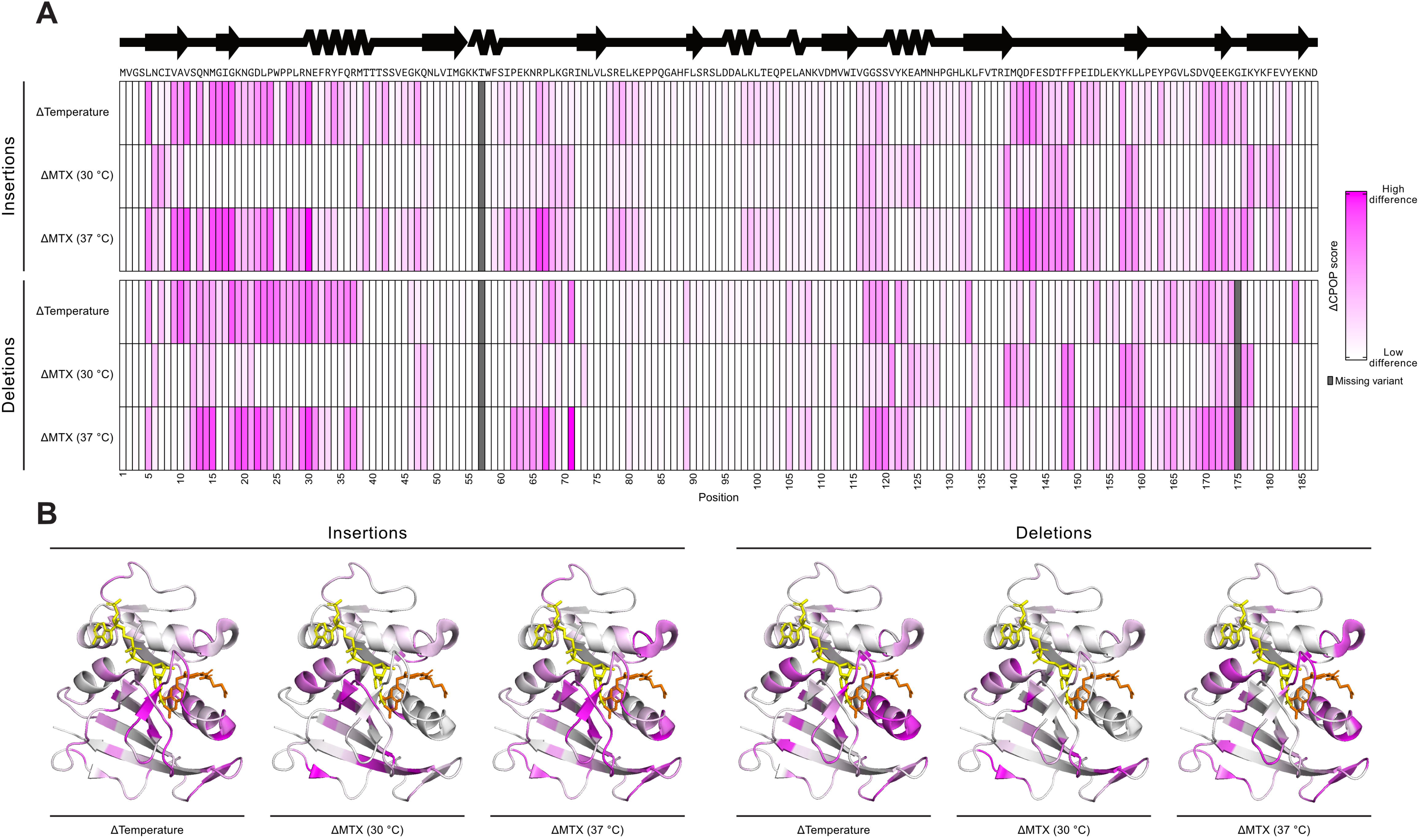
The effects of temperature and MTX on folding of the DHFR indel variants. (A) Heatmaps of ΔTemperature and ΔMTX CPOP scores for insertions and deletions at the four different conditions for each position in DHFR. The ΔTemperature scores were calculated by subtracting the score at 37 °C from the score at 30 °C. The ΔMTX scores were calculated by subtracting the score without MTX from the score with MTX. The ΔCPOP scores range from low difference (white) to high difference (magenta). Missing variants are colored grey. The amino acid sequence and a linear representation of the secondary structure of human DHFR are shown above the heatmap. (B) The ΔCPOP scores for insertions and deletions at the four different conditions are mapped to the DHFR structure (PDB: 1U72). The color scheme is the same as in panel A. NADPH is shown in yellow and MTX is shown in orange.

### Protein stability and AlphaFold2 predictions capture indel sensitivity

Next, we asked whether the experimentally determined indel effects in the CPOP system could be predicted using computational methods. Many computational tools for variant effect predictions have only been benchmarked for effects of substitution variants, and only return variant effect predictions for substitutions and not indels. However, a few variant effect predictors also predict indel effects, but with limited performance [27].

To overcome this challenge, we pursued a recently described approach that combines AlphaFold2 (AF2) and Rosetta to predict indel effects [9]. This strategy has recently proven effective in separating tolerated from non-tolerated deletion variants of a small α-helical protein. Specifically, we used AF2 to predict structures of DHFR indel variants tested in the CPOP system. This included all possible single amino acid deletions and single glycine insertions. For each indel variant, the AF2 output was five predicted models, of which we averaged the per-position pLDDT score across the entire protein. The pLDDT score, ranging from 0 to 100, reflects the confidence of AF2 in the predicted structure [43]. To obtain a relative metric of the accuracy of each indel structure prediction, ΔpLDDT scores were calculated per position by subtracting the wild-type pLDDT from the indel variant pLDDT. To predict the changes in folding stability (ΔΔG) between the wild-type and each indel variant, we used the five predicted indel AF2 models as input for Rosetta. We generated ten models per input structure (50 models in total), and the three lowest-energy scoring models were averaged to produce ΔGs for the wild-type and each indel variant. ΔΔG and ΔpLDDT correlated for both insertion and deletion variants and showed that indels predicted to have reduced structural stability by Rosetta also had lower confidence predictions by AF2 (**Fig. 6A**). Indeed, when colored by the experimentally obtained CPOP score, it was evident that detrimental indel variants tended to cluster in the upper-left corner, while wild-type-like indels were closer to a ΔΔG and ΔpLDDT of 0 (**Fig. 6A**). We noted that the range of ΔpLDDT scores was wider for insertions than deletions, indicating that AF2 was less confident about the structures of some predicted insertion variants compared to deletion variants (**Supplementary Fig. 8**). To assess the ability of ΔΔG and ΔpLDDT to distinguish detrimental indels from wild-type-like indels, we used receiver operating characteristic (ROC) analyses for each experimental condition and calculated the area under the curve (AUC) (**Fig. 6B**). To produce the ROC curves, we filtered the CPOP dataset to include only unique protein-level indel variants. This was done by removing insertion variants before glycine residues and removing one of the deletion variants in positions with two consecutive identical residues. Based on the distribution of CPOP scores (**Fig. 3A**), we selected a cutoff of 0.5 to categorize indels as detrimental or wild-type-like. In general, ΔΔG and ΔpLDDT predictions were better for deletions than for insertions (**Fig. 6B**). For both insertions and deletions, ΔpLDDT was the best predictor of the CPOP score at 37 °C with an AUC of 0.75 and 0.82, respectively (**Fig. 6B**). Predictions were most accurate for CPOP scores at 37 °C, perhaps because this is the condition with the largest dynamic range. However, this trend is only observed for ΔΔG for insertions (**Fig. 6B**). Interestingly, and particularly evident for insertions, ΔpLDDT outcompetes ΔΔG for most conditions, suggesting that AF2 pLDDT captures structural folding defects of indels better than Rosetta ΔΔG (**Fig. 6B**). Taken together, these results suggest that the combination of AF2 ΔpLDDT and Rosetta ΔΔG serves as a good indel effect predictor and can identify positions that are sensitive to indels.

**Fig. 6.**
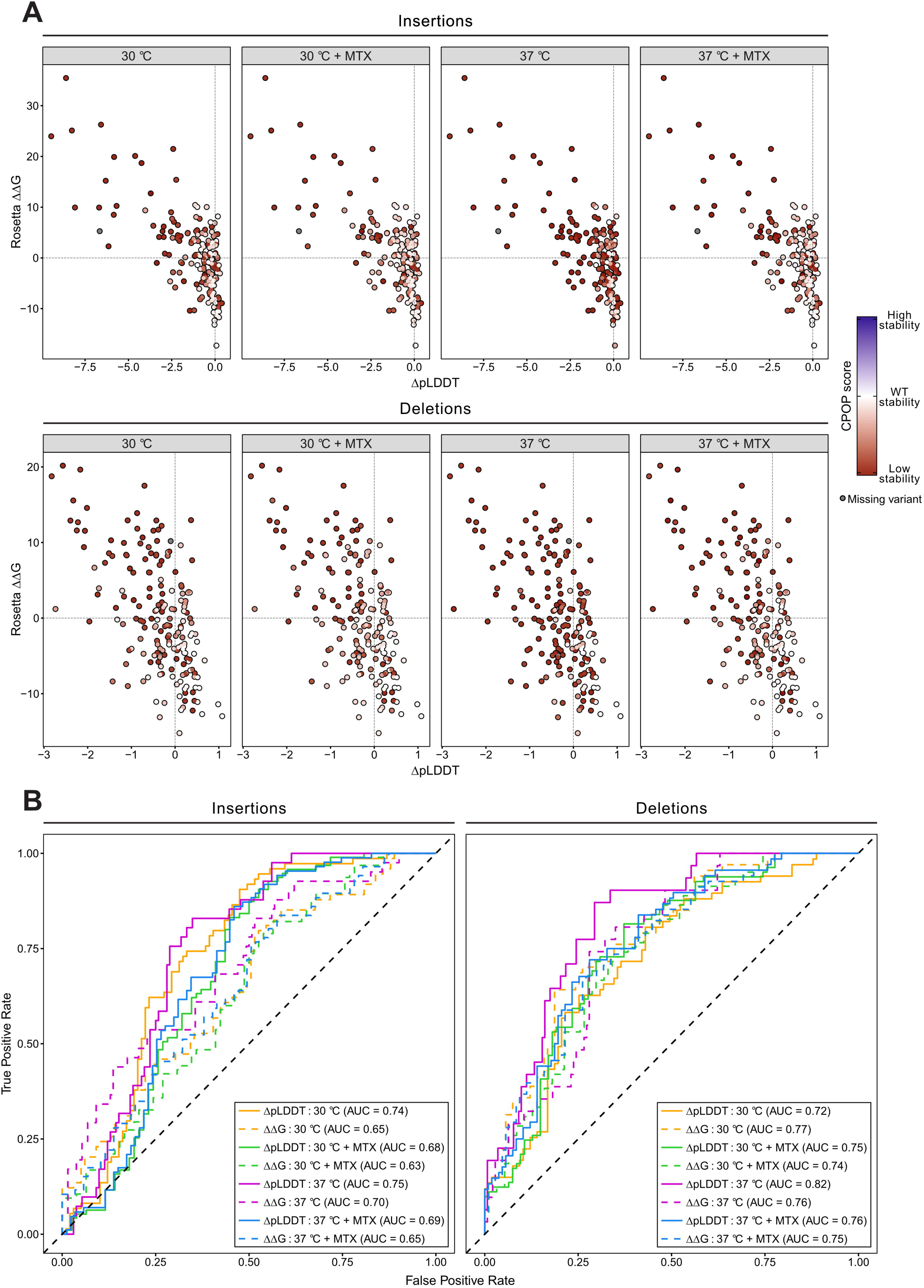
Predicting indel tolerance using Rosetta and AlphaFold2. (A) Rosetta ΔΔG (in Rosetta Energy Units) plotted against ΔpLDDT for each indel variant at the four different conditions. Indel variants are colored by the CPOP score and the color scheme is the same as in Fig. 3B. The dashed line on the y-axis and x-axis corresponds to the predicted wild-type stability and pLDDT, respectively. Positive ΔΔG scores indicate indel variants with reduced structural stability and negative ΔpLDDT scores indicate indel variants with reduced confidence in the predicted structure by AF2. (B) ROC curves for Rosetta ΔΔG (dashed lines) and AF2 ΔpLDDT (solid lines) to assess how well each measurement performs in separating detrimental from wild-type-like indel variants at the four different conditions. A cutoff of 0.5 was applied to the CPOP score. Indel variants with a CPOP score below 0.5 were categorized as detrimental, while those with a score above 0.5 were categorized as wild-type-like. The AUC for each predictor is reported in the legend. The dashed diagonal line denotes a random classifier.

## Discussion

Over the past few years, we have seen a substantial rise in large-scale experiments that probe mutational effects of missense variants [2, 25, 44–48]. These experiments have expanded our understanding of protein function and enabled us to understand the underlying mechanisms of several monogenic diseases. However, in comparison, indels have received much less attention and accordingly our current understanding of indel effects is greatly limited. Indels represent a class of variation that is dramatically different from side-chain changes introduced by substitutions. Altering the length of a protein affects the possible conformational space, and thus indels can have more substantial effects on protein function compared to missense variants. So far only a few reports have attempted to systematically address indels by DMS technologies [20–27]. This has resulted in a few general trends, such as proteins appearing to best tolerate indels in flexible loops as well as towards the N- and C-termini. However, since the indel tolerance varies widely both within and between proteins [27], we clearly have much more to learn about how indels affect protein structure and function.

Our results using the CPOP folding sensor on DHFR indels support many previous observations. For example, indels in DHFR seem best tolerated in the termini and internal loops, as with other proteins. However, the central loops spanning residues 60-70 and 75-88 are highly sensitive to indels, although both AF2 and Rosetta predict these residues to tolerate indels (**Supplementary Fig. 9**). One could speculate that the central loops, despite being exposed and flexible, are important for the registry of the neighboring β-sheets and binding of the co-factor NADPH which might support protein folding. In addition, we note that the α-helix at position 30-40 does allow deletions and these experimental findings are also supported by computational modelling (**Supplementary Fig. 9**). In general, we find that insertions are tolerated better than deletions. This is in agreement with observations on other proteins [24, 26], and likely reflects that it is easier to structurally accommodate an insertion, e.g. by extending a loop or forming a bulge, than handling the increased strain caused by a deletion [8]. For the central β-sheets, we observe a clear tendency for indel tolerance towards the ends of each strand. Also, particularly deletions are more detrimental in β-sheets compared to α-helices. These observations are in line with previous studies [24, 26].

An advantage of using DHFR as a model protein is that it binds tightly and specifically to MTX which results in an increased structural stability [31, 39]. This widens the dynamic range of the CPOP folding assay and allows us to identify highly destabilizing variants. Conversely, comparing the CPOP readout at 30 °C and 37 °C allows us to probe mildly destabilizing indels. Surprisingly, this revealed that 23% of the insertions and 24% of the deletions are temperature-sensitive (displaying a ΔTemperature score >0.4), and these indels cluster towards the N- and C-termini. Moreover, since most of the temperature-sensitive indels can be stabilized by MTX, this indicates that it is largely the increased dynamics conferred by high temperature that causes the structural instability and not an underlying cellular response (such as induction of molecular chaperones *etc*.). It is important to consider that our CPOP assay is optimized to measure protein folding stability, and hence, we cannot determine if DHFR indel variants are compromised in substrate- or co-factor binding or catalysis. Such detailed mechanistic insights would require complementary assays to disentangle indel effects on different properties important for DHFR function.

In recent years, we have seen tremendous progress in our ability to accurately predict protein structures from amino acid sequences. The development of AF2 [49] has provided predicted structures for essentially all known proteins. In parallel, other computational tools have been developed to predict the effects of missense variants [3, 4], resulting in powerful tools for both clinical genetics and protein engineering. However, due to sparse datasets, it still remains challenging to develop and validate prediction methods for how indels affect protein function. To build a predictive model of indel effects it appears important to consider sequence position and secondary structure of the mutation. Indeed, average substitution effects per position in combination with sequence position have provided relatively accurate estimates of indel effects in other recent studies [27].

We used a combination of AF2 and Rosetta, which has previously shown promising results for systematic deletions in a small protein domain [9], to separate tolerated from detrimental indels in DHFR. Our data shows that AF2 ΔpLDDT provides relatively accurate predictions of deletion effects on protein folding of DHFR (AUC = 0.82). For both insertions and deletions, Rosetta ΔΔG predicts folding defects slightly worse than ΔpLDDT. This suggests that Rosetta ΔΔG struggles to capture the larger-scale structural and conformational rearrangements required to accommodate indel variations. Additionally, for our CPOP assay, ΔpLDDT performs better in predicting indel effects on DHFR than two variant effect predictors (CADD and PROVEAN) and a large language model (ESM1b) do at predicting indel effects for several diverse protein domains, as measured by *in vitro* thermodynamic folding stability [25, 27]. Considering the performance of AF2 in predicting indel effects, it will be interesting to see in the future how effectively AF2 or its successors can predict effects of clinically relevant indels and be utilized for protein engineering.

## Materials and Methods

### Plasmids and cloning

The DNA sequences of the empty CPOP construct [30] and wild-type *E. coli* OPRTase (UniProt ID: P0A7E3) were codon-optimized for yeast expression and cloned into pDONR221 (Genscript). The DNA sequences of human DHFR (UniProt ID: P00374) and *E. coli* TrxA (UniProt ID: P0AA25) were codon-optimized for yeast expression and inserted into the CPOP construct and cloned into pDONR221 (Genscript). Missense variants were created by Genscript. For expression in yeast cells, entry clones were cloned into pAG415GPD-ccdB (Addgene plasmid #14146; http://n2t.net/addgene:14146; RRID:Addgene_14146, [50]) using Gateway cloning (Invitrogen).

### Yeast strains and media

Yeast strains were obtained from the Euroscarf collection. The wild-type yeast strain was BY4741. The *ura5*Δ*ura10*Δ yeast strain was created by first converting the *ura10::G418* strain to *ura10::NAT* by allele replacement [51]. Then the *ura10::NAT* cassette was PCR amplified and transformed into the *ura5::G418* strain. Small scale yeast transformations were performed using lithium acetate [52]. Untransformed yeast cells were cultured in yeast extract, peptone, dextrose (YPD) medium (2% glucose, 2% tryptone, 1% yeast extract), and transformed yeast cells were cultured in synthetic complete (SC) medium (2% glucose, 0.67% yeast nitrogen base without amino acids (Sigma), 0.2% synthetic drop-out supplement (Sigma)). Solid media was prepared using 2% agar.

### Yeast growth assays

For growth assays on solid media, yeast cells were cultured overnight (30 °C) to exponential phase. The cultures were harvested and washed twice in sterile water (3000 g, 5 minutes) and resuspended in sterile water. The cultures were adjusted to an OD_600nm_ of 0.4 and used for five-fold dilution series in sterile water. Then, cultures were spotted onto agar plates using 5 μL of each dilution. The plates were briefly air-dried before they were incubated at the indicated temperature for 2-3 days. Solid media with uracil was prepared using synthetic drop-out supplement without leucine, and solid media without uracil was prepared using synthetic drop-out supplement without leucine, uracil, and tryptophan, which was supplemented with 76 mg/L tryptophan. Plates containing MTX were prepared using 220 μM MTX (Sigma-Aldrich) and 1 mM sulfanilamide (Sigma-Aldrich).

### Library design

Full-length human *DHFR* was divided into five tiles of 39 or 40 residues that overlapped by three residues. The residues included in each tile were: tile 1: 1-40, tile 2: 38-77, tile 3: 75-114, tile 4: 112-151 and tile 5: 149-187. The library was synthesized and cloned into pAG415GPD-ccdB by Twist Bioscience and delivered in 96-well plates. Each well of the 96-well plates contained four variants for a single position: 1) a deletion of the residue at the position, 2) a single glycine (codon: GGT) insertion after the position, 3) a synonymous wild-type (silent) variant at the position and 4) a stop codon variant (nonsense mutation) at the position. All programmed DHFR variants were included in the libraries, except for those at position 57, which failed in library preparation by Twist Bioscience. Each well also contained the wild-type sequence. Wells were appropriately pooled to construct the desired five tiles. From here on, libraries in each tile were treated separately. First, 2 μL of each tile was electroporated at 1.8 kV into 50 μL NEB 10-beta electrocompetent *E. coli* cells (New England Biolabs). Immediately after electroporation, cells were suspended in 948 μL NEB 10-beta stable outgrowth medium and incubated for 1 hour (37 °C) to recover. Then, 100 μL of the transformation culture was diluted and plated on LB agar with ampicillin and incubated overnight (37 °C) for counting the number of single colonies. The remaining 900 μL of the transformation culture was transferred to 200 mL LB medium with ampicillin and incubated overnight (37 °C). At least 20,000 colonies were counted for each tile. The following day, plasmid DNA was isolated and purified from cultures using the NucleoBond Xtra Midi kit (Macherey–Nagel).

### Library transformation and selection

The DHFR indel libraries were transformed in parallel into the *ura5*Δ*ura10*Δ strain using lithium acetate and a 30-fold scale-up as described in [53]. Briefly, yeast cells were cultured overnight (30 °C) to late-exponential phase. The cultures were diluted in at least 150 mL YPD to an OD_600nm_ of 0.3 and incubated until the cells had completed two divisions. The cultures were harvested and washed three times in sterile water (3000 g, 5 minutes, room temperature) and the pelleted cells were resuspended in a transformation mix composed of: 7.2 mL 50% PEG-3350 (Sigma), 1.08 mL 1 M LiAc (Sigma), 300 μL 10 mg/mL freshly denatured salmon sperm DNA (Sigma), 1.2 mL sterile water and 30 μg library plasmid DNA diluted in 1.02 mL sterile water. The cells were gently suspended by inversion and incubated in a water bath (40 minutes, 42 °C). The cells were harvested (3000 g, 5 minutes, room temperature) and resuspended in 30 mL sterile water. Then, 5 μL of the transformation culture was plated on SC-leucine agar in duplicate and incubated for two days (30 °C) for counting the number of single colonies. The remaining volume of the transformation culture was transferred to a sufficient volume of SC-leucine medium to reach an OD_600nm_ of 0.2 and incubated for two days (30 °C). At least 20,000 colonies were counted for each tile. After the transformation culture reached saturation, 9 OD_600nm_ units were harvested in triplicates (17,000 g, 1 minute, room temperature) and the cell pellets were stored at −20 °C. These samples served as the control condition prior to selection. For the four selection conditions, 4×5 OD_600nm_ units were harvested in triplicates and washed three times in sterile water (17,000 g, 1 minute, room temperature) and resuspended in 400 μL sterile water. Then, 100 μL were plated on square plates (144 cm^2^) for each condition: 1) SC-uracil at 30 °C, 2) SC-uracil with MTX at 30 °C, 3) SC-uracil at 37 °C, 4) SC-uracil with MTX at 37 °C. Plates containing MTX were prepared using 220 μM MTX (Sigma-Aldrich) and 1 mM sulfanilamide (Sigma-Aldrich). Plates were briefly air-dried before they were incubated at 30 °C or 37 °C for two days. After incubation, lawns of colonies had formed on all plates and yeast cells were scraped off the plates using 10 mL sterile water. From each plate, 9 OD_600nm_ units were harvested (17,000 g, 1 minute, room temperature) and the cell pellets were stored at −20 °C. These samples served as the selection conditions. Finally, plasmid DNA was isolated from the pelleted cells using the ChargeSwitch Plasmid Yeast Mini kit (Invitrogen) and DNA concentrations were measured using the Qubit 2.0 Fluorometer (Invitrogen) with the Qubit dsDNA HS Assay Kit (Thermo Fisher Scientific).

### Amplicon preparation and library sequencing

Amplicons for each tile were prepared from the isolated plasmid DNA using specific primers with Illumina adapter sequences. For each tile, 50 µL PCRs were prepared composed of: 5 ng isolated plasmid DNA, 1× Phusion High-Fidelity PCR Master Mix with HF Buffer (New England Biolabs), 0.5 µM forward primer, 0.5 µM reverse primer and nuclease-free water (New England Biolabs) up to a total volume of 50 µL. The PCRs were subjected to the following program: initial denaturation at 98 °C for 30 seconds, followed by 7 cycles of denaturation at 98 °C for 15 seconds, annealing at 59-61 °C for 30 seconds, extension at 72 °C for 20 seconds, followed by a final extension at 72 °C for 2 minutes and 4 °C hold. The oligonucleotide sequences of the forward and reverse primers with Illumina adapter sequences and annealing temperatures for each tile are listed in the supplementary material (Supplemental File 1). After the first round of PCR, 15 µL of the product was mixed thoroughly with 15.75 µL Ampure XP beads (Beckman Coulter) (1.05× bead/PCR product ratio) and incubated at room temperature for 5 minutes. Then, the tubes were placed on a magnetic rack to pellet the amplicons bound to the beads and the supernatant was aspirated. The beads were washed twice with 200 µL 70% (v/v) ethanol and after the last wash, residual ethanol was completely removed by air-drying the beads at room temperature for 15 minutes. The amplicons were eluted with 16 µL nuclease-free water (New England Biolabs) and vortexed thoroughly. The tubes were pulse-centrifuged and placed on a magnetic rack to precipitate the magnetic beads separated from the cleaned-up amplicons. Then, 15 µL of the supernatant containing the amplicons was transferred to fresh PCR tubes to serve as template for another round of PCR to add Illumina index sequences and cluster-generating sequences onto the amplicons. For the second round of PCR, 50 µL PCRs were prepared composed of: 15 µL cleaned-up amplicons from the first round of PCR, 1× Phusion High-Fidelity PCR Master Mix with HF Buffer (New England Biolabs), 0.5 µM i5 index primer, 0.5 µM i7 index primer and nuclease-free water (New England Biolabs) up to a total volume of 50 µL. The PCRs were subjected to the following program: initial denaturation at 98 °C for 30 seconds, followed by 12 cycles of denaturation at 98 °C for 15 seconds, annealing at 65 °C for 30 seconds, extension at 72 °C for 20 seconds, followed by a final extension at 72 °C for 2 minutes and 4 °C hold. After the second round of PCR, 15 µL of the product was mixed thoroughly with 12 µL Ampure XP beads (Beckman Coulter) (0.8× bead/PCR product ratio) and incubated at room temperature for 5 minutes. The subsequent steps were performed as described for the clean-up of the product after the first round of PCR. The concentration of the final amplicons was measured using the Qubit 2.0 Fluorometer (Invitrogen) with the Qubit dsDNA HS Assay kit (Thermo Fisher Scientific), and the size and integrity of the final amplicons were evaluated using the 2100 Bioanalyzer (Agilent Technologies) with the High Sensitivity DNA kit. Finally, the concentration of amplicons was adjusted and all samples from all tiles were pooled in one tube. The pooled amplicons were subjected to paired-end next-generation sequencing using the Illumina NextSeq 550 System with the Mid Output v2.5 300 cycle kit (Illumina).

### Analysis of sequencing data

Paired-end reads in the raw FASTQ files were merged using FLASH2 [54]. FLASH2 was run with default parameters, but with --max-overlap=999 to disregard maximal overlap length. Then, adapters were trimmed using cutadapt [55], which was run with default parameters. Finally, the merged and trimmed FASTQ files were processed using an awk one-liner to count unique reads and write their DNA sequences and counts to a new TXT file:

`awk ‘NR%4==2{print}’ trimmed_merged.fastq | sort | uniq -c | sort -g > /trimmed_merged_counts.txt`.

To call DHFR variants and calculate CPOP scores, we wrote a Python script (https://github.com/KULL-Centre/_2024_Larsen-Ledet_CPOP/blob/main/functions.py) that used the awk-generated TXT files as input. First, sequences shorter or longer than the expected length of deletion or insertion variants, respectively, were removed. Then, each sequence was assigned one mutation type (insertion, deletion, synonymous, or nonsense) along with the position of the mutation. An additional filter was applied to remove out-of-frame sequences and sequences with more than one mutation at the protein level. For indel variants, sequences with a synonymous mutation at the protein level in addition to the programmed indel mutation were tolerated. Finally, a CPOP score was calculated for each DHFR variant within each tile.

For each replicate *n* (*n*=3) and each condition *t* (no selection, 30 °C, 30 °C + MTX, 37 °C and 37 °C + MTX), the count of each variant *v* was normalized to the sum of all synonymous variants *syn*:

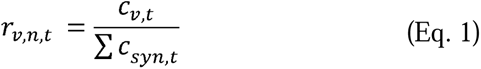

For each variant *v* and condition *t*, the three ratios *r_v,1,t_*, *r_v,2,t_*, and *r_v,3,t_* (one for each replicate) were averaged to produce a single ratio *r’_v,t_* for each condition:

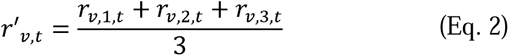

The standard deviation *s* of *r’_v,t_* was calculated with normalization performed using *n – 1:*

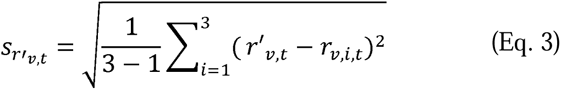

For each variant, the selection ratios *r’_v,sel_* for the four selection conditions (30 °C, 30 °C + MTX, 37 °C and 37 °C + MTX) were normalized to the control ratio *r’_v,ctrl_* for the control condition (no selection) to produce a score ratio *R_v,sel_* for each selection condition:

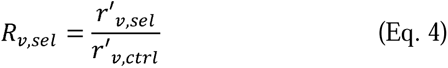

The standard deviation *s* of *R_v,sel_* was calculated using the formula for propagation of uncertainty for division:

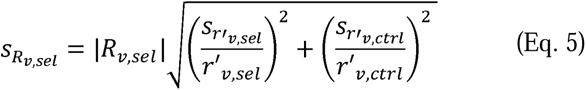

For each variant, the natural logarithm was taken of the score ratio R_v,sel_ to produce a final CPOP score for each selection condition:

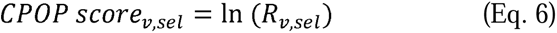

The final standard deviation *s* of *CPOP score_v,sel_* was calculated:

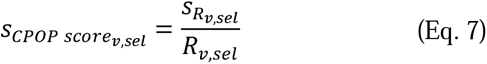

The final CPOP scores *CPOP score’_v,sel_* were rescaled to range from 0 to 1, where 0 represents the average score of nonsense variants and 1 represents the average score of synonymous variants. This ensures that CPOP scores across all five tiles are comparable and interpretable within the same scale. Importantly, the nonsense and synonymous variants exhibited a tight distribution around 0 and 1, respectively (**Supplementary Table 1**).

The final standard deviation *s’* of *CPOP score’_v,sel_* was rescaled in the same way by dividing it by the difference between the highest and lowest CPOP score prior to rescaling:

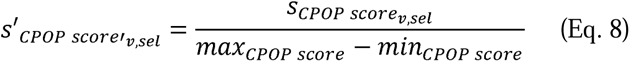

For technical reasons related to sequencing, the following replicates were excluded: tile 2, replicate 2, 30 °C; tile 2, replicate 2, 37 °C + MTX; tile 3, replicate 1, 37 °C.

### Structural analysis

The monomeric CPOP-DHFR fusion protein structure was predicted by AF2 and aligned to each monomer of the crystal structure of *E. coli* OPRTase (PDB: 1ORO) [56]. AF2 was run with ColabFold [57]. Specifically, the full amino acid sequence of the CPOP-DHFR fusion protein was used as query sequence and the notebook was run with default parameters. The five generated models had similar pLDDT scores. To produce the CPOP-DHFR homodimer, one of the CPOP-DHFR monomeric models was selected and duplicated in PyMOL using the copy command. Then, each of the CPOP-DHFR monomers was aligned to PDB: 1ORO using the align command in PyMOL. We tried to predict the CPOP-DHFR homodimer using AF2-Multimer [58]. However, the N- and C-termini of CPOP were poorly predicted with low pLDDT scores in the homodimer compared to in the monomer (**Supplementary Fig. 10**).

SSDraw [59] was used to produce the linear representation of the secondary structure of DHFR. The AF2 predicted structure of wild-type DHFR was used as PDB entry.

The software was run using the Google Colab notebook at: https://colab.research.google.com/github/ethanchen1301/SSDraw/blob/main/SSDraw.ipynb.

The secondary structure elements were assigned for DHFR (PDB: 1U72) [60] using DSSP. DSSP assigns the secondary structure for each residue based on the atomic coordinates of the input structure. DSSP uses seven different secondary structure categories, which we grouped into three broader groups: loops, helices, and strands. Loops include “bends”, “turns” and “none”, helices include “α-helices”, “310-helices” and “π-helices”, and strands include “β-strands” and “β-bridges”.

The relative solvent accessible surface area (rSASA) was calculated for DHFR (PDB: 1U72) using DSSP.

The distances to the MTX binding site were calculated between heavy atoms of MTX and the C_β_ atoms of the DHFR residues, with C_α_ used for glycine.

### Modelling of wild-type DHFR and indel variants

AF2 was used to predict structural models of human wild-type DHFR and indel variants. Modelling of the wild-type structure was based on PDB: 1U72. From the wild-type sequence (UniProt ID: P00374), FASTA files were produced for each DHFR indel variant by mutating the wild-type amino acid sequence to all possible single amino acid deletions and a single glycine insertion after each residue. AF2 (version 2.2.4) was run with the following flags: --max_template_date=2021-11-01 and --run_relax=False. AF2 outputs five models of the predicted structure for both the wild-type and each indel variant. The pLDDT scores were calculated as the average per-position pLDDT from the five output models. The ΔpLDDT scores were obtained by subtracting the wild-type pLDDT score from the DHFR indel variant pLDDT score.

### Structural stability predictions

The Rosetta relax protocol was used to predict the changes in thermodynamic stability of indel variants. Rosetta relax was run on each of the five output structures predicted by AF2 for both the wild-type and each indel variant. The Rosetta relax output was ten models per input structure, resulting in a total of 50 models for both the wild-type and each indel variant. The three lowest-energy scoring models were averaged to produce a single ΔG of folding for both the wild-type and each indel variant. The ΔΔG values were calculated by subtracting the wild-type ΔG value from the DHFR indel variant ΔG value, accounting for the change in amino acid length due to a deletion (−1) or an insertion (+1):

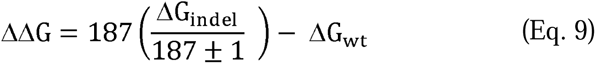

Rosetta relax (version 2021, August, c7009b3) was run with the following flags: -nstruct=10 and -relax:constrain_relax_to_start_coords=1.

## Acknowledgements

We acknowledge the use of the sequencing and computing core facilities at the Department of Biology, University of Copenhagen. We thank Anne-Marie Lauridsen for technical assistance and Vasileios Voutsinos for assistance with Illumina sequencing. Fig. 2D was created with BioRender.com.

## Supplemental files

This article includes the following supplemental information:

- Supplemental figures (SupplementalFigures.pdf).
- Supplemental dataset (SupplementalFile1.xlsx).

## Data and code availability

All data generated are included in the figures and supplemental files. Sequencing reads are available at the NCBI Gene Expression Omnibus (GEO) repository (accession number: GSE270811). The code is available at GitHub (https://github.com/KULL-Centre/_2024_Larsen-Ledet_CPOP).

## Author contributions

S.L.-L., S.L., A.P., S.G., C.H.S., A.V.G., H.L., and S.V.N performed the experiments. S.L.-L., A.P., S.G., A.V.G., H.L., S.V.N., J.R.W., A.S. and R.H.-P. analyzed the data. K.L.-L., J.R.W. and R.H.-P. conceived the study. S.L.-L. and R.H.-P. wrote the paper.

