## Supplemental Figures for "Systematic characterization of indel variants using a yeast-based protein folding sensor"

### *Supplementary Information*

**Fig. S1**, *Vector map of pAG415GPD-CPOP-DHFR.* *p.2*

**Fig. S2**, *Sequencing counts correlate between replicates across all tiles and conditions.* *p.3*

**Fig. S3,** *The standard deviation is low for most indel variants.* *p.4*

**Fig. S4**, *Low-throughput validation of selected indel variants.* *p.5*

**Fig. S5**, *Deletions are generally more detrimental than insertions.* *p.6*

**Fig. S6**, *DHFR indels show no correlation with the rSASA.* *p.7*

**Fig. S7**, *The ΔMTX scores exhibit low correlation with the distance to the MTX binding site.* *p.8*

**Fig. S8**, *AF2 predicts insertion variants with slightly less confidence than deletion variants.* *p.9*

**Fig. S9**, *Computational modelling captures some positions sensitive to indels.* *p.10*

**Fig. S10**, *AF2 predicts the CPOP-DHFR homodimer with low confidence.* *p.11*

**Table S1**, *Nonsense and synonymous variants are centered around 0 and 1, respectively.* *p.12*

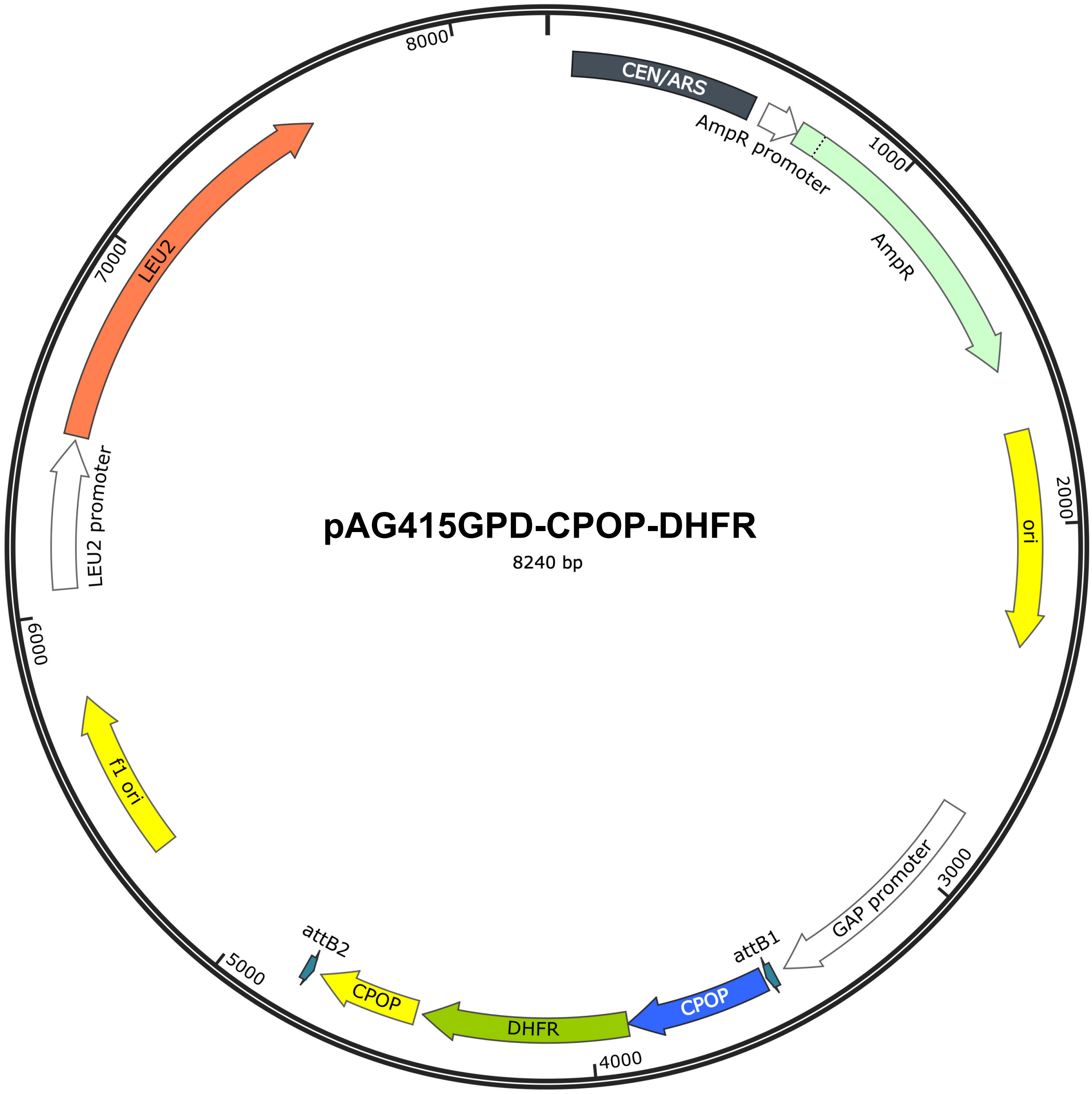


**Fig. S1.** *Vector map of pAG415GPD-CPOP-DHFR.* Vector map of the CEN-based expression vector pAG415GPD with a *LEU2* marker. The CPOP-DHFR fusion protein was expressed from the constitutive GAP promoter. The SnapGene® software (from Dotmatics; available at snapgene.com) was used for generating the plasmid map.

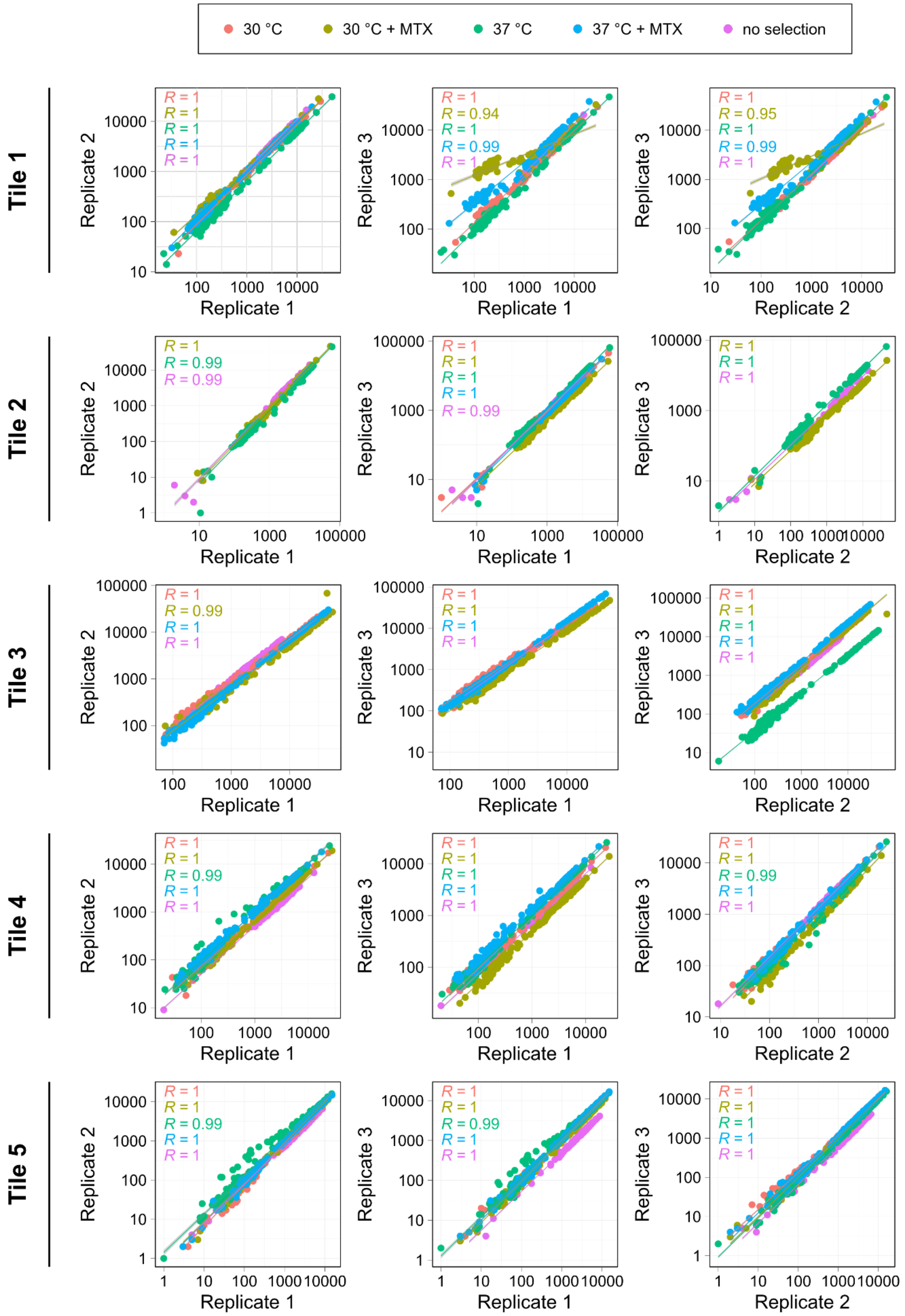


**Fig. S2.** *Sequencing counts correlate between replicates across all tiles and conditions.* Plots showing the count correlation for all DHFR variants across three replicates for each tile, colored by condition (no selection, 30 °C, 30 °C + MTX, 37 °C and 37 °C + MTX). The Pearson correlation coefficient (R) is provided for each condition.

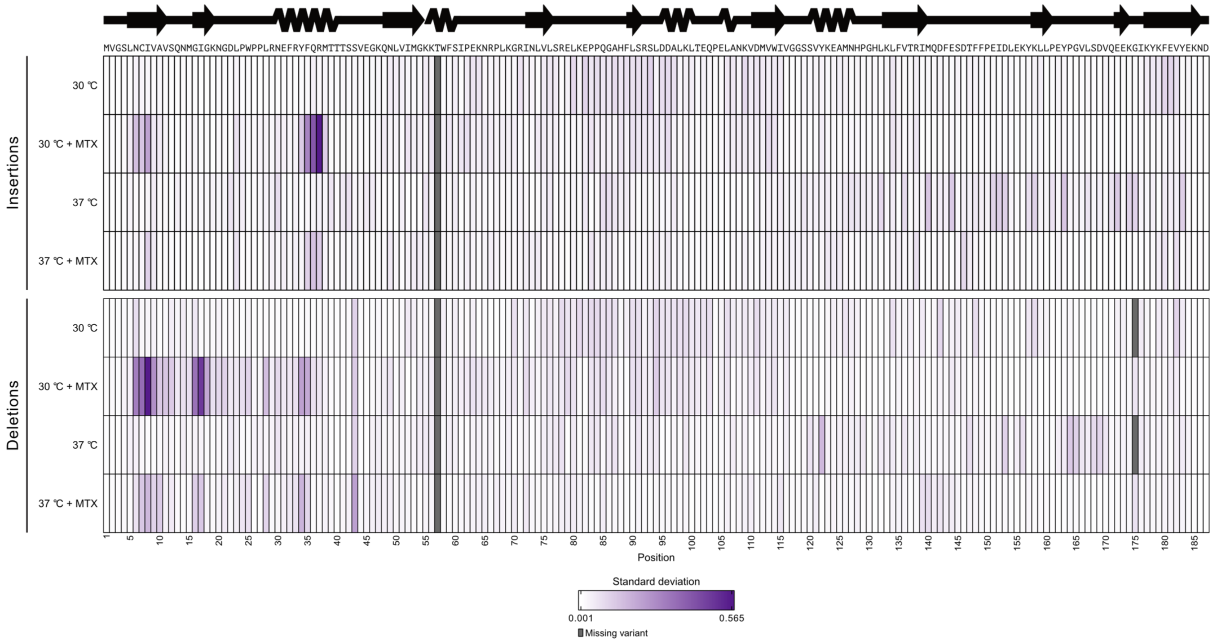


**Fig. S3.** *The standard deviation is low for most indel variants.* Heatmaps of the standard deviation of CPOP scores at the four different conditions. The standard deviations range from the lowest deviation (white) to the highest deviation (purple). Missing variants are colored grey. The amino acid sequence and a linear representation of the secondary structure of human DHFR are shown above the heatmap.

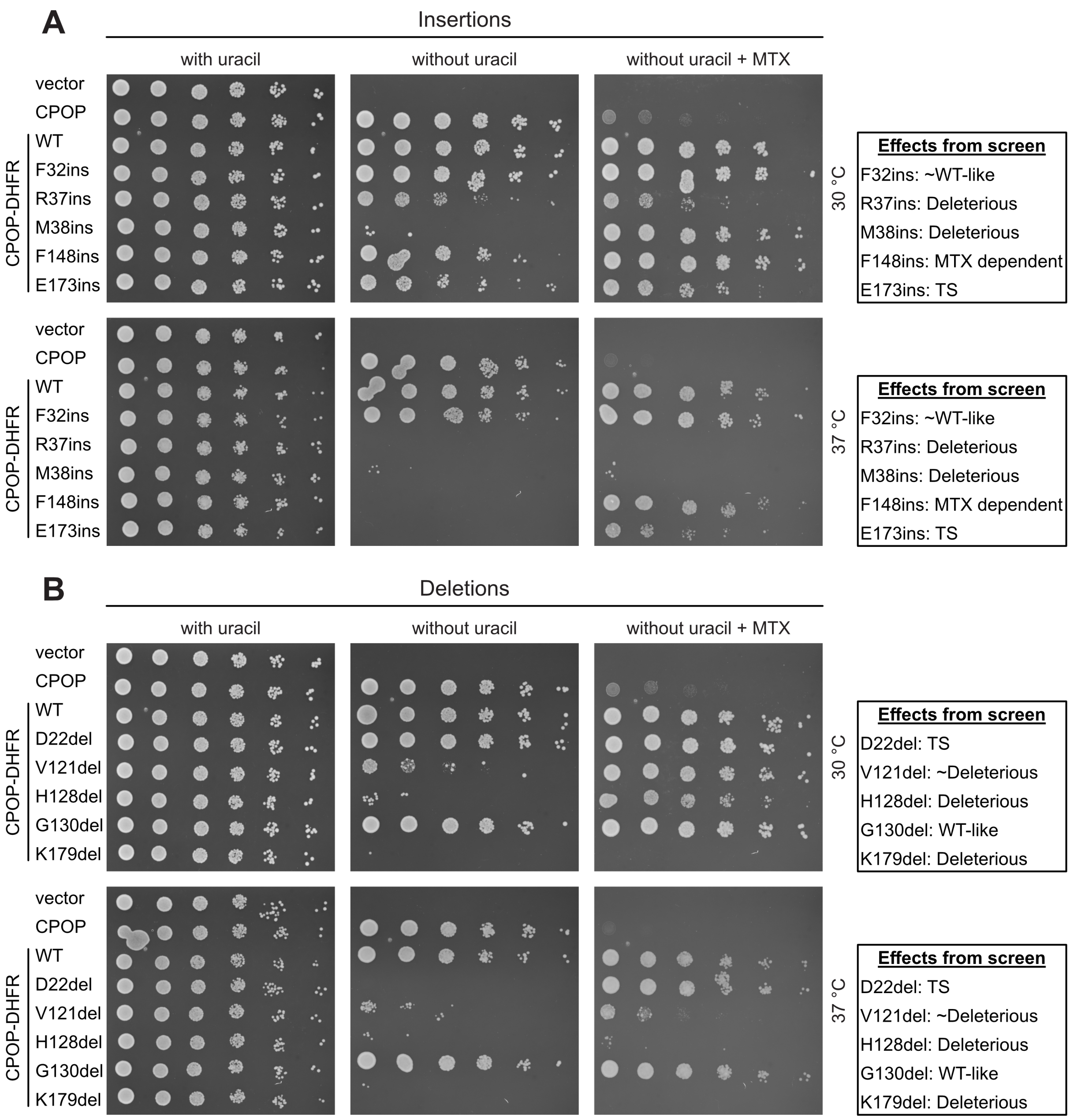


**Fig. S4.** *Low-throughput validation of selected indel variants.* (A) Comparison of growth of the *ura5*Δ *ura10*Δ strain transformed with either an empty vector, CPOP or CPOP fusion constructs with wild-type human DHFR or an insertion variant. Cells were grown at 30 °C and 37 °C on medium with and without uracil, and without uracil with MTX. (B) Same as in (A), but for deletion variants.

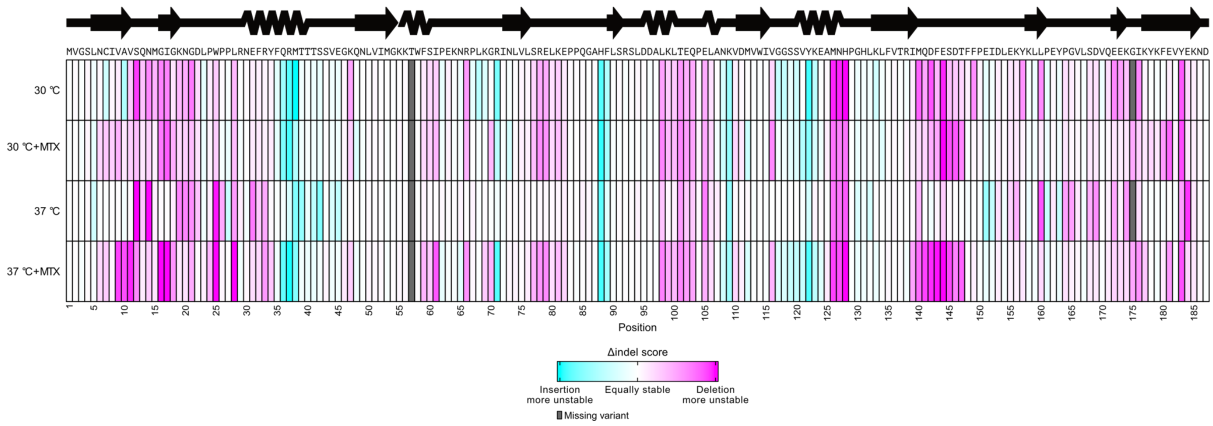


**Fig. S5.** *Deletions are generally more detrimental than insertions*. Heatmaps of Δindel CPOP scores at the four different conditions. The Δindel scores were calculated by subtracting the deletion score from the insertion score. The Δindel scores range from insertions being more unstable (cyan), over insertions and deletions being equally stable (white), to deletions being more unstable (magenta). Missing variants are colored grey. The amino acid sequence and a linear representation of the secondary structure of human DHFR are shown above the heatmap.

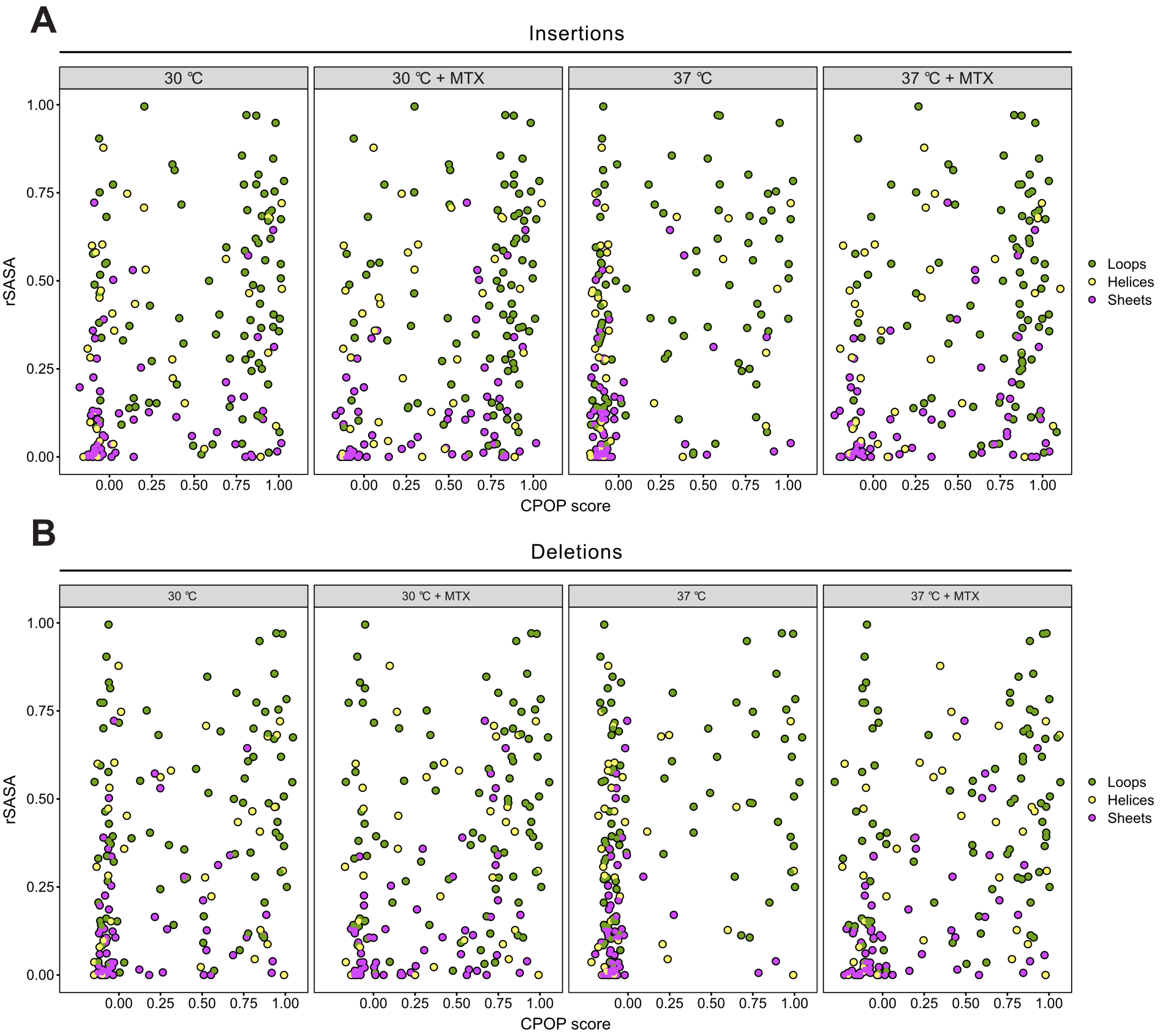


**Fig. S6.** *DHFR indels show no correlation with the relative solvent accessible surface area (rSASA).* (A) rSASA plotted against the CPOP score for each insertion variant at the four different conditions. Residues are colored by the secondary structural element. A rSASA value of 1 indicates that the residue is completely exposed, while 0 denotes completely buried. (B) Same as in (A), but for deletion variants.

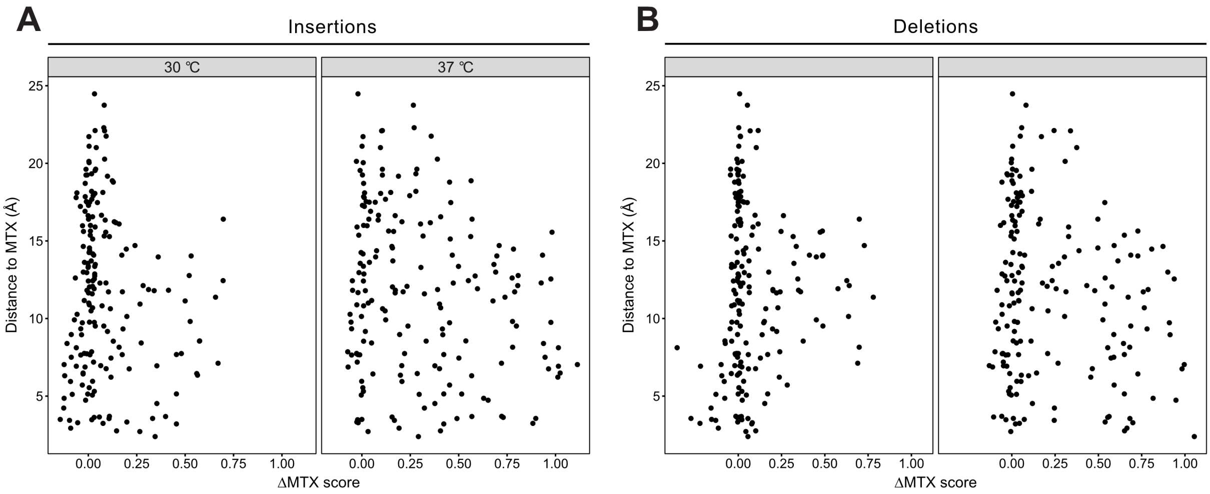


**Fig. S7.** *The ΔMTX CPOP scores exhibit low correlation with the distance to the MTX binding site.* (A) The distance of each residue to the MTX binding site in Å plotted against the ΔMTX CPOP scores for each insertion variant at the four different conditions. Higher ΔMTX CPOP scores indicate a larger stabilizing effect of MTX. (B) Same as in (A), but for deletion variants.

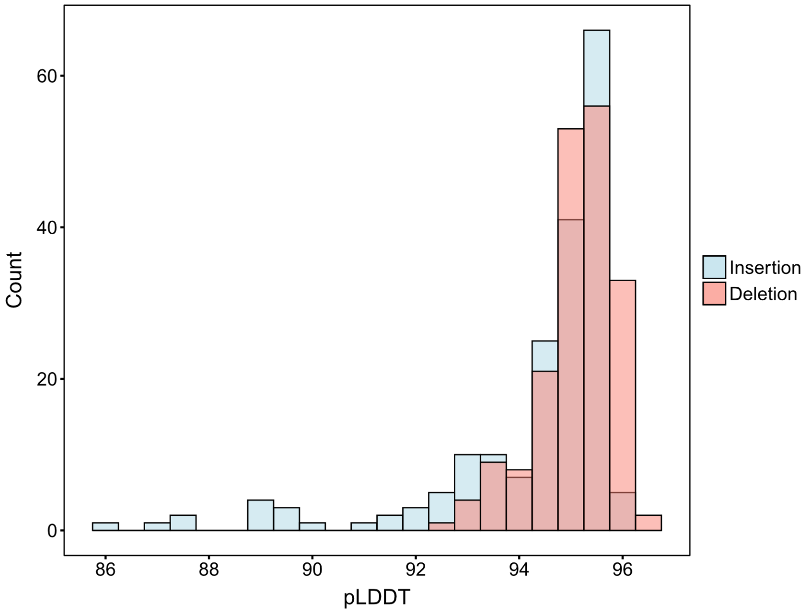


**Fig. S8.** *AF2 predicts insertion variants with slightly less confidence than deletion variants.* Histograms of AlphaFold2 (AF2) average pLDDT across all positions for each predicted indel structure. Higher pLDDT scores indicate more confidence by AF2 in the predicted structure.

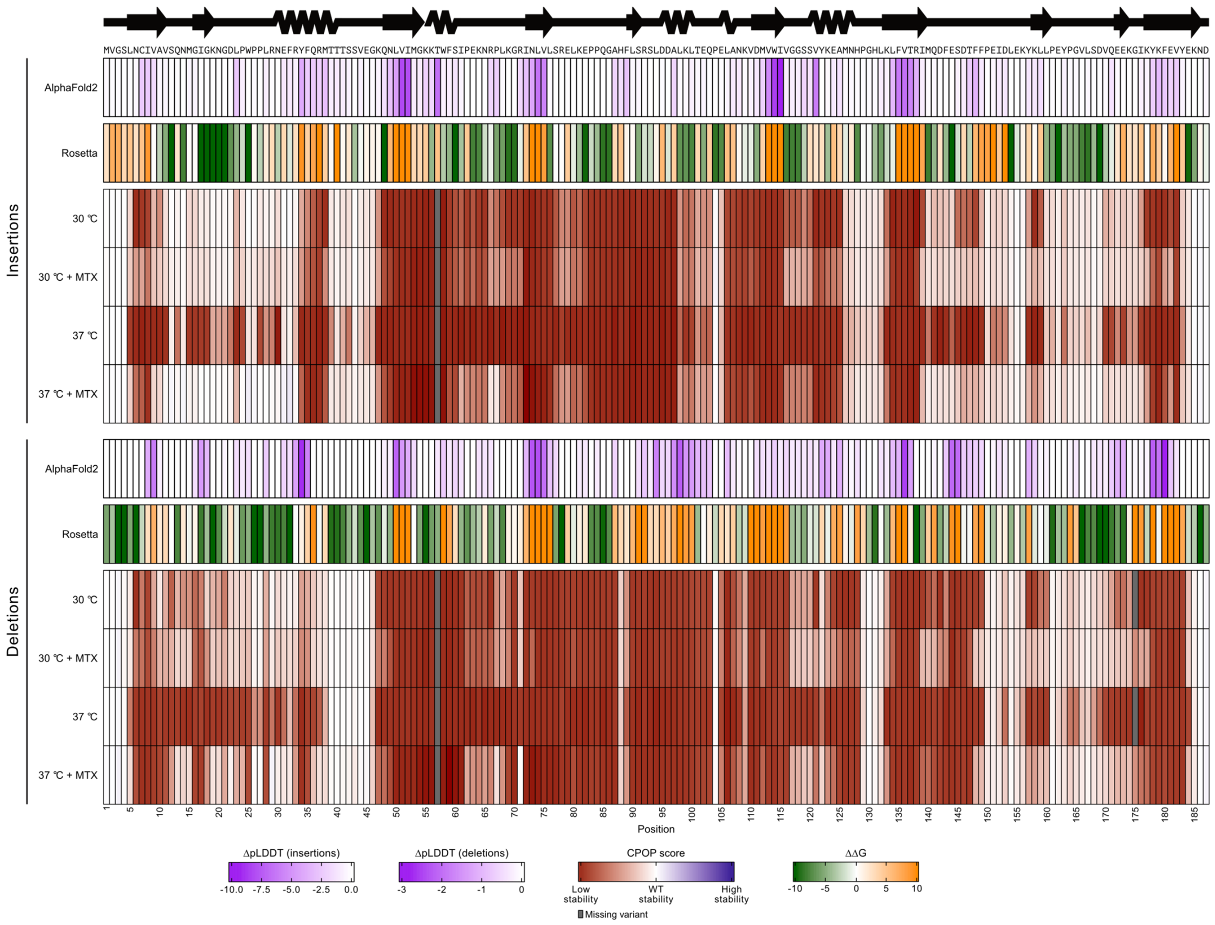


**Fig. S9.** *Computational modelling captures some positions sensitive to indels.* (A) Heatmaps of the experimental CPOP scores at the four different conditions for insertions and deletions. Above each heatmap, heatmaps of AlphaFold2 ΔpLDDT and Rosetta ΔΔG are included. Both ΔpLDDT and ΔΔG predict the α-helix at position 30-40 to be more intolerant to insertions than deletions. The central loops spanning residues 60-70 and 75-88 are not predicted to be sensitive to indels.

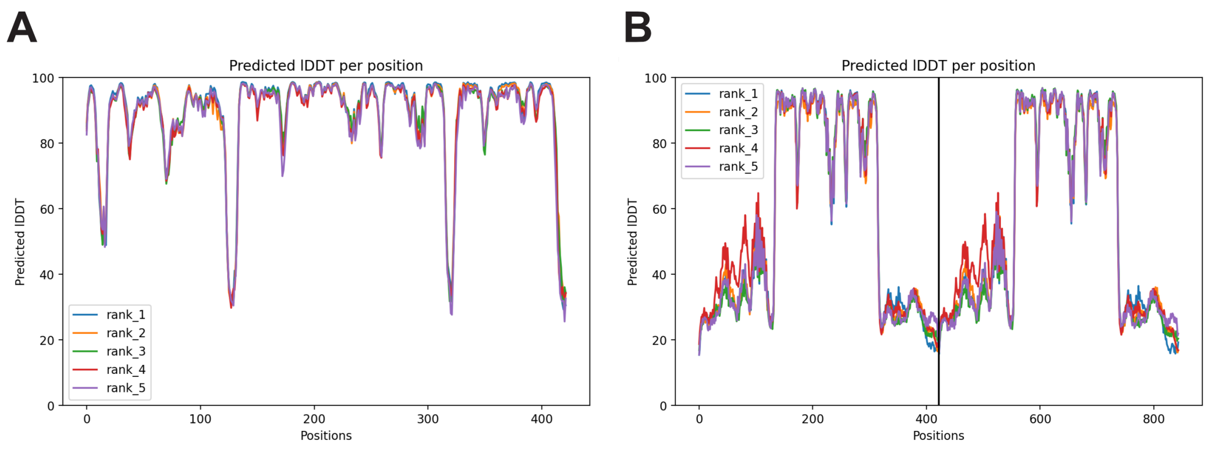


**Fig. S10.** *AF2 predicts the CPOP-DHFR homodimer with low confidence.* (A) pLDDT per residue for the CPOP-DHFR monomer. The N-terminus of CPOP is position 1-123, DHFR is position 131-317 and the C-terminus of CPOP is position 326-425. The low pLDDT scores at position 124-130 and 318-325 are linkers. (B) pLDDT per residue for the CPOP-DHFR homodimer. The positions are defined as in (A). Clearly, the N- and C-termini of CPOP are predicted with less confidence in the homodimer as compared to the monomer in (A).

| **Nonsense variants (μ = 0)** | | |
|---|---|---|
| | **2σ** | **Within μ ± 2σ** |
| **30 °C** | 0.06 | 97% |
| **30 °C + MTX** | 0.07 | 97% |
| **37 °C** | 0.06 | 95% |
| **37 °C + MTX** | 0.06 | 95% |

| **Synonymous variants (μ = 1)** | | |
|---|---|---|
| | **2σ** | **Within μ ± 2σ** |
| **30 °C** | 0.05 | 96% |
| **30 °C + MTX** | 0.05 | 96% |
| **37 °C** | 0.04 | 96% |
| **37 °C + MTX** | 0.07 | 98% |

**Table S1.** *Nonsense and synonymous variants are centered around 0 and 1, respectively.* Most nonsense variants have CPOP scores close to 0, while most synonymous variants have CPOP scores close to 1. The mean (μ) is defined as 0 for nonsense variants and 1 for synonymous variants. The fractions of variants within two standard deviations (2σ) from the mean are calculated.
